# Cortical silencing results in paradoxical fMRI overconnectivity

**DOI:** 10.1101/2020.08.05.237958

**Authors:** Carola Canella, Federico Rocchi, Shahryar Noei, Daniel Gutierrez-Barragan, Ludovico Coletta, Alberto Galbusera, Stefano Vassanelli, Massimo Pasqualetti, Giuliano Iurilli, Stefano Panzeri, Alessandro Gozzi

**Affiliations:** Functional Neuroimaging Laboratory, Center for Neuroscience and Cognitive systems, Istituto Italiano di Tecnologia, Rovereto, Italy; Center for Mind and Brain Sciences, University of Trento, Rovereto, Italy; Neural Computational Laboratory, Center for Neuroscience and Cognitive Systems, Istituto Italiano di Tecnologia, Rovereto, Italy; University of Padua, Italy; Biology Department, University of Pisa, Pisa, Italy; Systems Neurobiology Laboratory, Center for Neuroscience and Cognitive systems, Istituto Italiano di Tecnologia, Rovereto, Italy

## Abstract

fMRI-based measurements of functional connectivity are commonly interpreted as an index of anatomical coupling and direct interareal communication. However, causal testing of this hypothesis has been lacking. Here we combine neural silencing, resting-state fMRI and *in vivo* electrophysiology to causally probe how inactivation of a cortical region affects brain-wide functional coupling. We find that chronic silencing of the prefrontal cortex (PFC) via overexpression of a potassium channel paradoxically increases rsfMRI connectivity between the silenced area and its thalamo-cortical terminals. Acute chemogenetic silencing of the PFC reproduces analogous patterns of overconnectivity, an effect associated with over-synchronous fMRI coupling between polymodal thalamic regions and widespread cortical districts. Notably, multielectrode recordings revealed that chemogenetic inactivation of the PFC attenuates gamma activity and increases delta power in the silenced area, resulting in robustly increased delta band coherence between functionally overconnected regions. The observation of enhanced rsfMRI coupling between chemogenetically silenced areas challenges prevailing interpretations of functional connectivity as a monotonic index of direct axonal communication, and points at a critical contribution of slow rhythm generators to the establishment of brain-wide functional coupling.

## Introduction

A rapidly expanding approach to understand neural organization is to map large-scale patterns of spontaneous neural activity in the resting brain via non-invasive neuroimaging. The ease and reproducibility of resting state fMRI (rsfMRI) have promoted the widespread use of this approach, leading to the observation that spatio-temporal patterns of spontaneous fMRI activity are organized into highly coherent functional networks, defined by temporally correlated fluctuations in BOLD signal ^1^. The non-invasive nature of rsfMRI has fueled the use of this method to map intrinsic brain network organization in the healthy human brain, as well as in psychiatric or neurological conditions, in which evidence of disrupted or aberrant rsfMRI functional coupling has been largely documented ^2, 3^. However, despite the growing popularity of rsfMRI, our knowledge of the underpinnings and neural drivers of brain-wide fMRI coupling remains very limited.

Statistical dependencies in spontaneous fMRI signal are commonly interpreted as an index of interareal functional communication and axonal connectivity^1^. This interpretation is supported by empirical and computational evidence suggesting that structural and rsfMRI-based connectivity are robustly related. For example, structural and functional connection strengths are correlated both at the whole-brain and mesoscopic scale^4-6^, and rsfMRI network topography closely recapitulates patterns of anatomical connectivity^7, 8^. In keeping with this, experimental resection of callosal connections^9^ and chemogenetic inactivation of the amygdala have been shown to reduce functional connectivity with regions anatomically linked to the manipulated areas^10^. Computational modelling supports this view, as synchronous rsfMRI fluctuations can be modelled by dynamical systems endowed with realistic anatomical connectivity patterns of long-range axonal interactions^11^, and simulated axonal lesions in these models result in reduced functional coupling^12^.

However, despite a prominent body of research supporting structurally-based interpretations of rsfMRI connectivity, evidence arguing against a dyadic relationship between structural and rsfMRI connectivity has also been accumulating. For example, robust rsfMRI coupling among brain regions not directly structurally connected has been reported in acallosal humans, primates and rodents ^9, 13, 14^. Similarly, neurological disorders such as Parkinson’s disease, multiple sclerosis or stroke are often associated with unexpectedly increased interareal rsfMRI connectivity, despite the focal or generalized loss of cortical function characterizing these conditions^15, 16^. These observations argue against a strictly monotonic relationship between structural and functional connectivity and call for a deeper investigation of the causal role of direct axonal connectivity in shaping large-scale functional coupling.

Perturbational approaches are critically required to disentangle the neural drivers of brain-wide rsfMRI coupling for two reasons. First, the correlative nature of rsfMRI makes measures of functional connectivity ambiguous indicators of direct interareal interaction, as synchronous signals could reflect not only direct interaction among areas, but also other sources of covariation, including the contribution of global or large-scale slow fMRI fluctuations ^17-19^, or the effect of indirect long-range loops. Second, targeted deconstructions of the fundamental elements governing rsfMRI coupling could uncover the cause-effect nature of patterns of rsfMRI dysconnectivity associated with brain pathologies, enabling a back-translation of dysconnectivity into interpretable neurophysiological events and models that can help understand, diagnose or treat brain disorders. This line of inquiry is of great importance, as prevalent clinical findings of increased connectivity upon loss of cortical function remain at present mechanistically unexplained^16^.

Here we combine rsfMRI, neural and chemogenetic silencing (chemo-fMRI^20^) and in vivo electrophysiology to probe how inactivation of cortical activity causally affects rsfMRI coupling. Surprisingly, we find that chronic and acute neural silencing of the mouse medial prefrontal cortex (PFC), a core component of the mouse default mode network DMN, ^21^ results in paradoxically increased rsfMRI coupling between the silenced area and its thalamo-cortical terminals, an effect associated with decreased gamma activity in the targeted regions, and increased delta band neural coherence between overconnected areas. Our data suggest that rsfMRI connectivity does not monotonically reflect direct axonal output under all brain states and conditions, and reveal a critical contribution of large-scale low-frequency fluctuations to the establishment of brain-wide functional coupling.

## Results

### Chronic silencing of the prefrontal cortex leads to rsfMRI overconnectivity

Structurally based models of rsfMRI connectivity predict that neural silencing of a network node would result in diminished functional coupling with regions receiving direct axonal projections from the affected substrate^10, 12^. To causally test this prediction, we carried out rsfMRI measurements in a cohort of mice in which neuronal activity in the medial prefrontal cortex (abbreviated here as PFC) was chronically inhibited via bilateral viral transduction of the inward rectifying potassium channel Kir2.1 under the control of a pan-neuronal promoter (Fig. S1A). Prior research has shown that viral-mediated Kir2.1 expression results in a reduction of both evoked and spontaneous neuronal excitability lasting several weeks ^22, 23^. In keeping with this, *in vivo* electrophysiological recordings in the PFC of mice unilaterally transfected with Kir2.1 revealed a robust reduction of spontaneous firing rate in the targeted cortical area with respect to its control contralateral region (N=4, paired t test, *p* = 0.002, Fig. S2).

We next compared the patterns of rsfMRI connectivity in Kir2.1 and GFP-transduced control littermates, by imaging Kir2.1 transfected mice four weeks after viral injections (Fig. 1A, Fig. S1A). Consistent with previous investigations ^24, 25^, seed-based probing revealed significant long-range correlation between the PFC and thalamo-cortical components of the mouse DMN in both cohorts (Fig. 1B). Surprisingly, between-group comparisons revealed foci of significantly *increased* rsfMRI connectivity in the posterior cingulate/retrosplenial cortex and centromedial thalamic regions of Kir2.1 transfected mice (t test, p < 0.05, *t* > 2.03, FWE cluster-corrected, p < 0.05; Fig. 1C). Regional quantifications of DMN connectivity via multiple prefrontal-DMN seeds corroborated these findings, revealing increased rsfMRI synchronization along the entire midline extension of this network (two-way ANOVA, F_(1, 33)_ = 6.93; p *=* 0.013; Fig. 1D) and its centromedial thalamic targets (t test, t_33_ = 2.589, p = 0.014; Fig. 1E). Voxel-wise mapping did not reveal any other foci of reduced functional connectivity (t > 2.03, cluster correction at p < 0.05). Importantly, all the thalamo-cortical regions showing increased rsfMRI connectivity in Kir2.1 mice are characterized by high projections density from the PFC, as seen by comparing the magnitude of inter-group rsfMRI connectivity differences with incoming axonal connectivity strength inferred from a voxel-model of the mouse brain connectome^6^ (Fig. 1F, Wilcoxon rank sum test, p<0.0001). Interestingly, the direction and the location of DMN rsfMRI overconnectivity was not altered by global fMRI signal regression (Figure S3), with the exception of thalamic areas, in which the effect was strongly attenuated. Together, these findings suggest that chronic inhibition of neural activity in the PFC paradoxically increases functional connectivity between long-range thalamo-cortical terminals of the mouse DMN.

**Figure 1.**
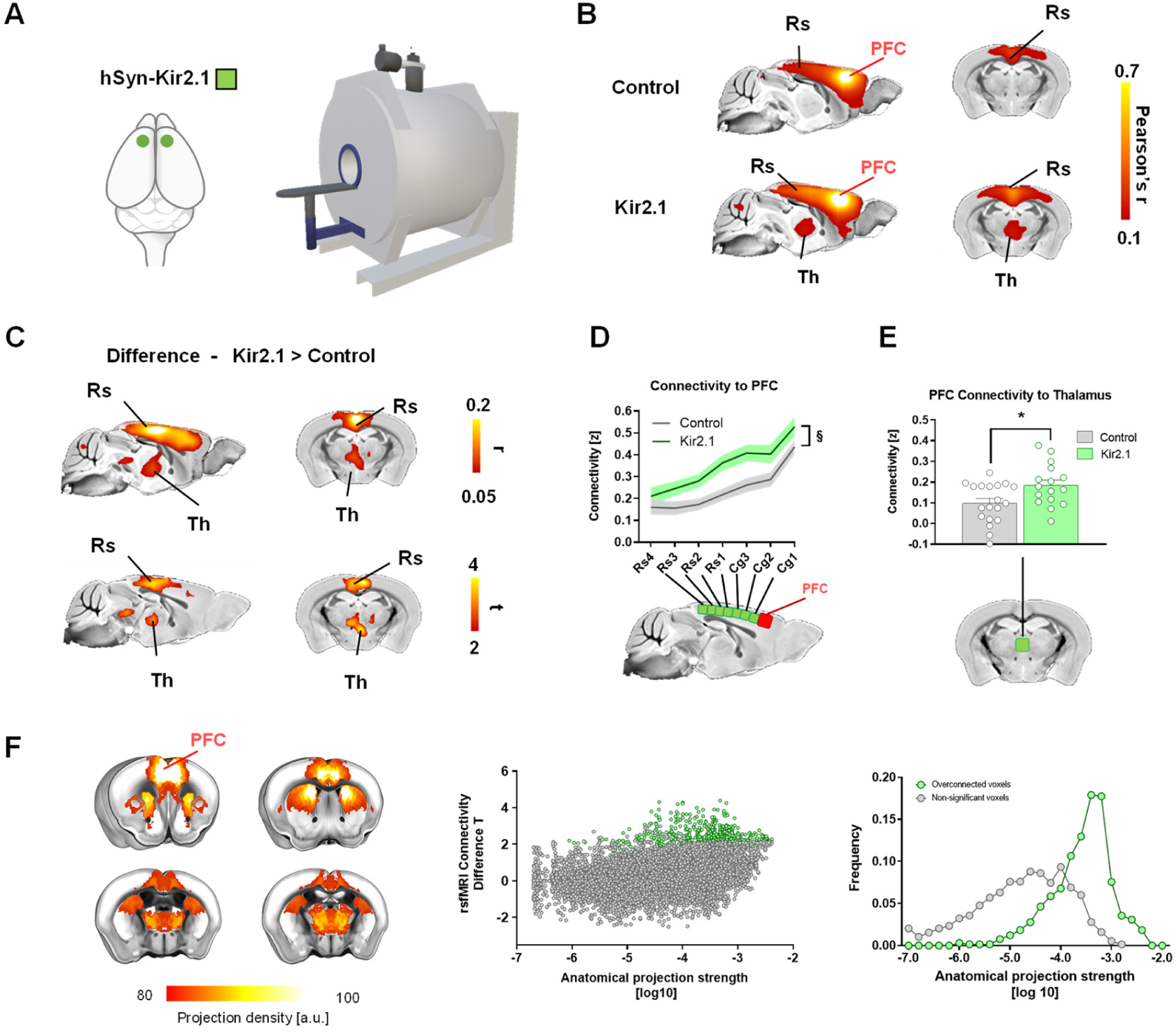
Chronic silencing of the PFC results in rsfMRI overconnectivity. **(A)** Summary of experimental design. The potassium channel Kir2.1 or GFP (control) were transduced bilaterally into the PFC of adult male mice. Four weeks after viral injections, mice underwent rsfMRI scanning. **(B)** Seed based connectivity mapping of the PFC in GFP (control), and Kir2.1-transduced subjects. **(C)** Corresponding group difference maps. Area with significantly increased rsfMRI connectivity in Kir2.1 expressing mice are depicted in red-yellow (r and T stat difference map). **(D)** Antero-posterior profiling of rsfMRI connectivity of the PFC within the midline axis of the mouse DMN. **(E)** Fronto-thalamic rsfMRI overconnectivity in Kir2.1 expressing mice. **(F)** Regions exhibiting rsfMRI overconnectivity in Kir2.1 mice are robustly innervated by the PFC. Left: Axonal projections from the PFC (top 20% strongest connections). Middle: scatter plot illustrating intergroup differences in rsfMRI connectivity as a function of PFC structural connectivity strength. Green dots indicate significantly functionally overconnected voxels. Right: Distribution of overconnected voxels as a function of axonal connectivity strength Green. DMN: Default Mode Network; Cg: cingulate cortex; PFC: prefrontal cortex, RS: retrosplenial; Th: Thalamus. § p<0.05, 2-way ANOVA, repeated measurements; * p<0.05, student t test.

### Chemogenetic inhibition of the prefrontal cortex leads to rsfMRI overconnectivity

The observation of increased DMN connectivity upon chronic inhibition of PFC activity argues against a tight dyadic relationship between axonal and rsfMRI-based functional connectivity. However, our results in Kir2.1 mice could reflect complex homeostatic or adaptive responses to chronic silencing. To corroborate the specificity of these findings and obtain mechanistic insight into the origin of the observed fMRI overconnectivity, we designed a new set of experiments in which DREADD-based chemogenetics was employed to induce a time-controlled, acute silencing of PCF activity during rsfMRI scanning. An overview of experimental procedures is provided in Figure 2A. To enable remote silencing of fronto-cortical activity, we bilaterally transfected the PFC with the inhibitory hM4Di DREADD using a pan-neuronal promoter (Figure 2A). Three weeks after viral injection, control (GFP-transfected) and hM4Di-expressing animals underwent rsfMRI scanning or electrophysiological recordings before and after intravenous injection of the DREADD activator clozapine-N-oxide (CNO). To account for the slow pharmacokinetic profile of CNO in the rodent brain ^26, 27^, both imaging and electrophysiological recordings were split into a pre-CNO injection baseline, a transitory (0-15 min) drug-equilibration period, and an active time window (15-50 min post CNO injection) to which all our analyses refer to, unless otherwise specified (Fig. 2B-C).

**Figure 2.**
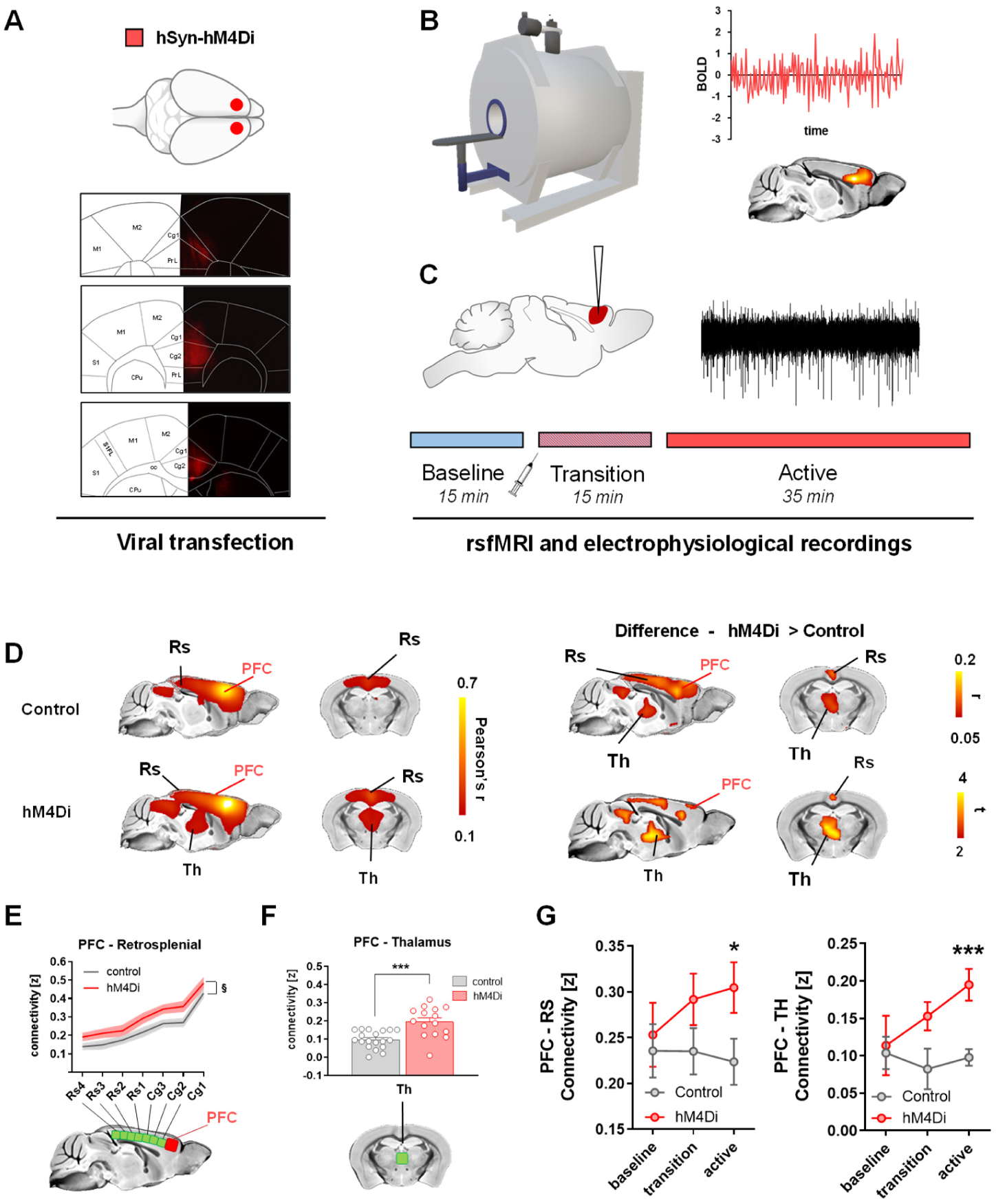
Chemogenetic silencing of the PFC results in rsfMRI overconnectivity. (A) Experimental design of chemo-fMRI experiments. AAV8-hSyn-hM4Di or AAV8-hSyn-GFP (control) were bilaterally injected into the PFC of wild type. The bottom panels show representative viral expression across three adjacent slices (viral injection in middle slice – see also Fig. S1B). Three weeks after transduction (B) mice underwent rsfMRI scanning or (C) electrophysiological recordings. A reference acquisition timeline is reported to depict timeseries binning into a 15 min pre-CNO reference baseline, a drug equilibration window (15 min, transition), and a 35 min CNO active time window (active). (D) Seed-based connectivity of the PFC and between group difference map revealed rsfMRI over-connectivity in the DMN of hM4Di expressing mice during the active phase. (E) Antero-posterior profiling of rsfMRI connectivity of the PFC along the midline axis of the mouse DMN in the two cohorts. (F) Thalamo-cortical rsfMRI hyper synchronization in hM4Di expressing mice. (G) Prefrontal-retrosplenial and nprefrontal-thalamic connectivity timecourse. Data are plotted as mean ± SEM. Cg: cingulate cortex; PFC: prefrontal cortex, RS: retrosplenial cortex; Th: Thalamus. § p<0.05, 2-way ANOVA repeated measurements, genotype effect; * p<0.05, ***<p<0.001, student t test.

To assess whether acute chemogenetic silencing of the PFC results in rsfMRI oversynchronization, we compared rsfMRI connectivity patterns in hM4Di transfected and control mice upon acute CNO administration (active phase, Fig. 2 and Fig. S1B). Recapitulating the Kir2.1 findings, voxel-wise mapping revealed foci of significantly increased rsfMRI connectivity in the posterior cingulate/retrosplenial cortices and midline thalamic regions of DREADD-expressing mice (t test, P < 0.05, FWE corrected, cluster defining threshold t > 2.03, p < 0.05; Fig. 2D). Regional quantifications corroborated the presence of rsfMRI oversynchronization along the cingulate and retrosplenial axis of the DMN, and between the PFC and medio-dorsal thalamic areas (2-way repeated measures ANOVA, F_1,32_ = 6.58; p = 0.0155; t test t_32_ = 4.30, p = 0.001, respectively; Fig. 2E-F), a set of regions characterized by dense incoming projections from the PFC (Figure S4, Wilcoxon rank sum test, p<0.0001). Notably, the direction and the anatomical location of the observed rsfMRI overconnectivity was regionally unaltered by global fMRI signal regression (Figure S5), arguing against an unspecific contribution of global fMRI co-activation or arousal related global dynamics^17, 18, 28^.

Moreover, baseline PFC connectivity in these areas was comparable across groups (voxel-wise mapping, Z>2.03 cluster corrected, PFC-Cingulate, 2 way ANOVA, F_1,32_ = 0.48, p = 0.49, Thalamo-PFC, t test, t_32_ = 0.23, p = 0.81), and oversynchronization gradually emerged in the hM4Di cohort after CNO administration, peaking during the DREADD active time-window (PFC-Rs: T_32_ = 2.158, p = 0.03, PFC-Th: T_32_ = 4.301, p = 0.0001, student t test, Fig. 2G). Moreover, no intergroup differences were observed in the characteristic hemodynamic response function in this area (kernel height p > 0.6; time-to-peak p > 0.12, full-width-at-half-peak p > 0.37, t test) nor were between-group differences in arterial blood pressure (p > 0.7, Student t test) or blood gas levels observed (P_a_CO_2_ p = 0.49; P_a_O_2_ p = 0.22, Student t-test). These control measurements rule out major spurious vascular or hemodynamic contributions and corroborate the specificity of the mapped changes. More broadly, our chemo-fMRI results show that acute inhibition of PFC activity results in a pattern of DMN oversynchronization closely recapitulating that observed with chronic Kir2.1-mediated silencing, suggesting that this phenomenon is not manipulation-specific, nor the indirect consequence of homeostatic reactivity to protracted neural silencing.

### Chemogenetic silencing of the prefrontal cortex increases interareal delta coherence

To probe the efficacy of chemogenetic silencing and investigate the neural correlates of the observed rsfMRI overconnectivity, we next carried out electrophysiological recordings in the PFC of hM4Di- or GFP-transduced control animals prior to and after CNO administration, under the same experimental conditions of rsfMRI (Fig. 3). Baseline electrophysiological traces revealed the presence of appreciable spontaneous multi-unit activity in the PFC both groups (mean firing rate 15.0 ± 2.2 spikes/s in hM4Di-expressing, and 14.1 ± 3.8 in GFP-transduced mice, n = 5 each group, p = 0.85, t test, Fig. 3AB). As expected, CNO administration robustly inhibited firing rate in hM4Di expressing mice, but not in control subjects (Fig. 3A-C, p<0.01 FDR corrected, t test). In keeping with the time-profile of rsfMRI overconnectivity, DREADD-induced PFC inhibition was characterized by gradual decrease of neural firing upon CNO administration, reaching a steady state approximately 15-20 min after the intravenous bolus (Fig. 3C).

**Figure 3.**
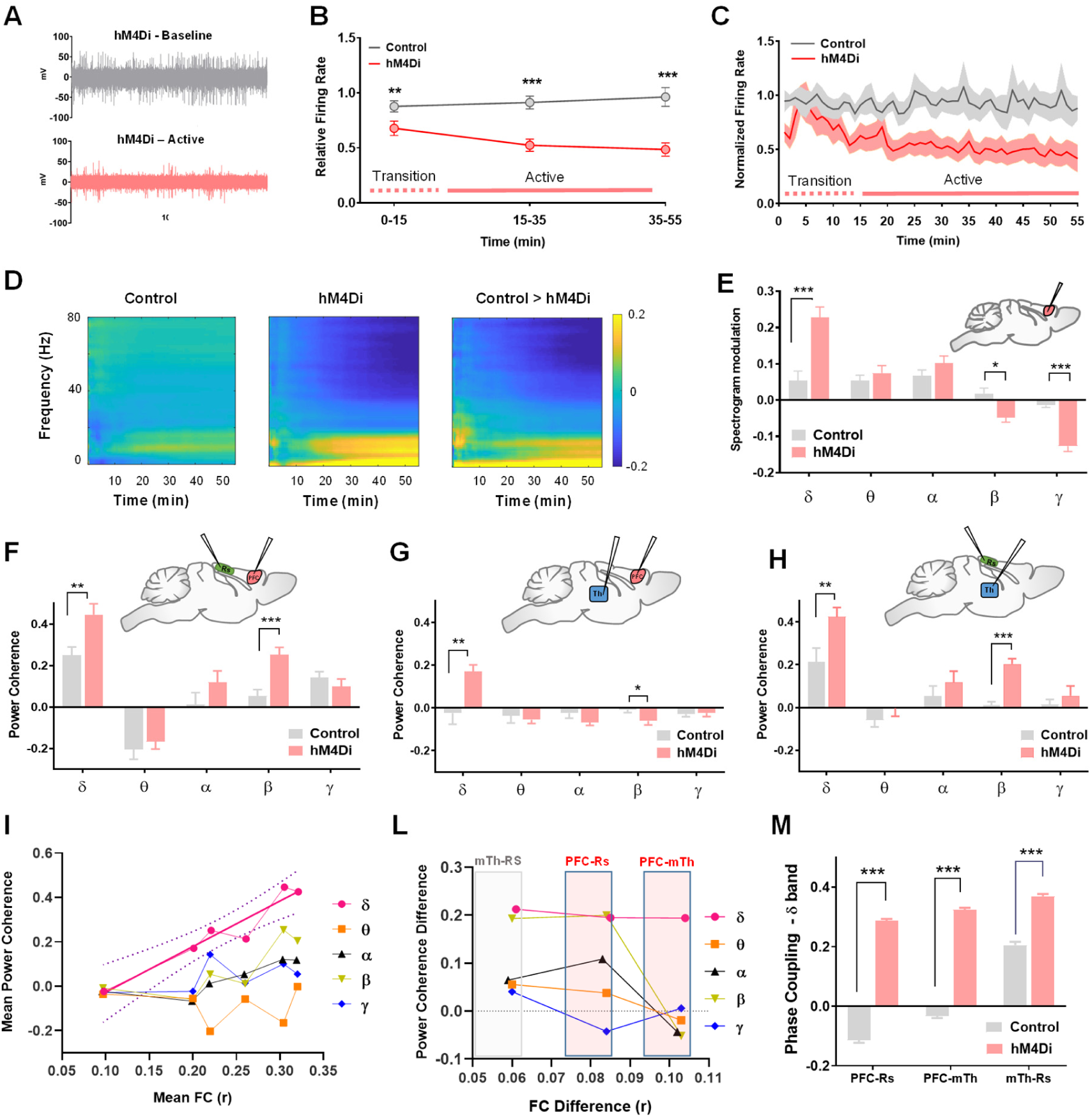
Chemogenetic silencing of the PFC results in decreased γ activity and increased interareal δ band coherence. (A) Representative raw traces collected before and after CNO injection in a representative recordings site of hM4Di animal. (B-C) Reduced firing rate in hM4Di-expressing mice compared to GFP-transduced controls. (D) Mean post-injection spectrogram in control (Left), hM4Di-expressing animals (center), and mean between group difference (right). (E) Quantification of band-specific power spectrum changes upon CNO injection in both groups (F-G-H). Baseline normalized power coherence at different frequency bands for PFC-Thalamus (Left), PFC-retrosplenial and Retrosplenial-thalamus electrode pairs (mean ±SEM; FDR corrected *q <0.05, **q<0.01, ***q<0.001, t test). (I) Correlation between mean band specific coherence and mean functional connectivity for all pairs of electrophysiologically-probed regions (PFC-Rs; PFC-Th; Rs-Th). Mean FC data were extracted for corresponding regional pairs (Fig. 2) during the CNO active time window in hM4Di and control animals. (L) Corresponding band specific coherence and mean functional connectivity difference (hM4Di – Control) for all pairs of electrophysiologically-probed regions in (I). (M) Baseline normalized phase coherence in delta band between Rs-thalamus electrode pair (mean ±SEM; *p <0.05, **p<0.01, ***p<0.001, t test, FDR corrected).

To obtain insight into the neural rhythms underlying the observed rsfMRI overconnectivity, we next analyzed local field potential (LFP) traces obtained from the same set of PFC electrophysiological recordings. Consistent with the observed firing rate reduction, LFP spectrograms from hM4Di-transduced animals revealed a robust decrease in β and γ power after CNO administration with respect to CNO treated controls (F_1,22_ = 239.4, p < 0.001, Fig. 3D). However, the power reduction in these frequency bands was also accompanied by a prominent increase in PFC δ power (defined here as 0.1 - 4 Hz band) in DREADD-expressing subjects (Fig. 3C-D, δ p < 0.001, θ p = 0.77; α p = 0.34; β p = 0.01; γ p<0.001; Student t test, Bonferroni corrected).

Slow rhythms such as δ band fluctuations are characterized by regionally extended synchrony that is at least in part mediated by a single higher order thalamic relay^29, 30^, whereas high-frequency oscillations such as α, β and γ rhythms are typically associated with more direct interareal communication between neural populations and cortical areas^31-33^. The observed increase in δ band power in the chemogenetically silenced PFC led us to hypothesize that the ensuing rsfMRI overconnectivity could be primarily driven by enhanced large-scale δ band coherence, rather than by a direct interareal communication via canonical higher frequency oscillations. To test this hypothesis, we carried out a set of simultaneous multi-electrode LFP recordings in the PFC, retrosplenial and centromedial thalamus of control and DREADD-transfected animals (Fig. 3 and S6). These regions were selected to cover the anatomical substrates exhibiting the highest functional overconnectivity in our prior chemo-fMRI studies (cf. Fig. 2). Spectral-power quantifications after CNO administration revealed a reduction in γ power in all the three recording sites, together with increased δ power in the PFC, but not in the retrosplenial cortex or centromedial thalamus of DREADD-transfected animals (Fig. S7). To further explore the possible neural correlates of fMRI connectivity in terms of frequency-dependent neural communication, following the logic of Wang et al.^34^, we next computed the synchronization between each of pair of regions separately for each frequency band, and compared the obtained coherence values with corresponding interareal rsfMRI coupling (Fig. 3F-M). Notably, only δ band power synchronization appeared to be fully concordant with our imaging findings, revealing robustly increased coherence between PFC-retrosplenial and PFC-thalamic electrode pairs. Interestingly, increased δ coherence was also observed between PFC and retrosplenial areas (Fig. 3F-H). Other bands did not show concordance with rsfMRI results: β band synchronization was increased after CNO administration in PFC-retrosplenial (and in retrosplenial-thalamus) electrode pairs, but was decreased in PFC-thalamus, while no significant change in power synchronization upon DREADD stimulation was observed in the θ and γ bands. (Fig. 3F-G). In keeping with this, we found that average pair-wise δ coherence across all the recorded sites exhibited a robust linear relationship with corresponding group-level pair-wise rsfMRI connectivity measured in both DREADD and control animals during the CNO active window (Fig. 3I, p=0.012, FDR corrected, R^2^ = 0.92). No significant relationship between coherence and pairwise rsfMRI connectivity was observed in any of the other bands (p > 0.11 all bands, FDR corrected). Similarly, we found no correlation between average band specific power (i.e. the mean power of each electrode pair) and pair-wise rsfMRI connectivity in any of the probed areas (p > 0.15, all bands, FDR corrected, Fig. S8), suggesting that rsfMRI connectivity is more plausibly explained by increased neural coherence, rather than the corresponding relative change in band specific power across electrode pairs. In keeping with this notion, a comparison of CNO-elicited band coherence differences with corresponding functional connectivity changes revealed that δ band was the only frequency range plausibly explaining the observed functional overconnectivity in PFC-retrosplenial and PFC-thalamic sites (Fig. 3L). These findings were paralleled by the observation of similarly robust increase in interareal δ phase coherence, a widely used measure of functional synchronization in LFP/EEG studies ^35^ (Fig. 3H). Taken together, these results corroborate a neural origin for our imaging findings, and implicate increased interareal coherence in the δ band as the most plausible driver of the observed rsfMRI over-synchronization.

### Chemogenetic silencing of the prefrontal cortex leads to functional oversynchronization between polymodal thalamus and widespread cortical areas

If δ band synchronization is a prominent neural correlate of the mapped overconnectivity, the observation of increased thalamo-retrosplenial δ coherence (Fig. 3H) implies that this pair of regions should similarly exhibit increased rsfMRI coupling. Such a scenario would be consistent with the presence of foci of chemo-fMRI overconnectivity in centro-medial and antero-dorsal areas of the polymodal thalamus, two PFC-innervated areas that serve as generators and cortical propagators of δ oscillatory rhythms, respectively ^29, 30^. The putative involvement of these polymodal thalamic areas as relays of slow oscillatory activity (and putatively fMRI overconnectivity) would in turn imply the possibility of spatially segregating thalamo-prefrontal rsfMRI overconnectivity profiles into a centro-medial (polymodal) and a lateral (unimodal) component, the former exhibiting rsfMRI over-synchronization with wider cortical territories.

To explore this hypothesis, we clustered voxels in the thalamus based on unique between-group functional connectivity differences ^36^. This approach led to the identification of two main thalamic sub-territories, one corresponding to the polymodal thalamus (including centromedial and anterodorsal areas), and the second encompassing latero-thalamic (sensory) areas (Fig. 4A).

**Figure 4.**
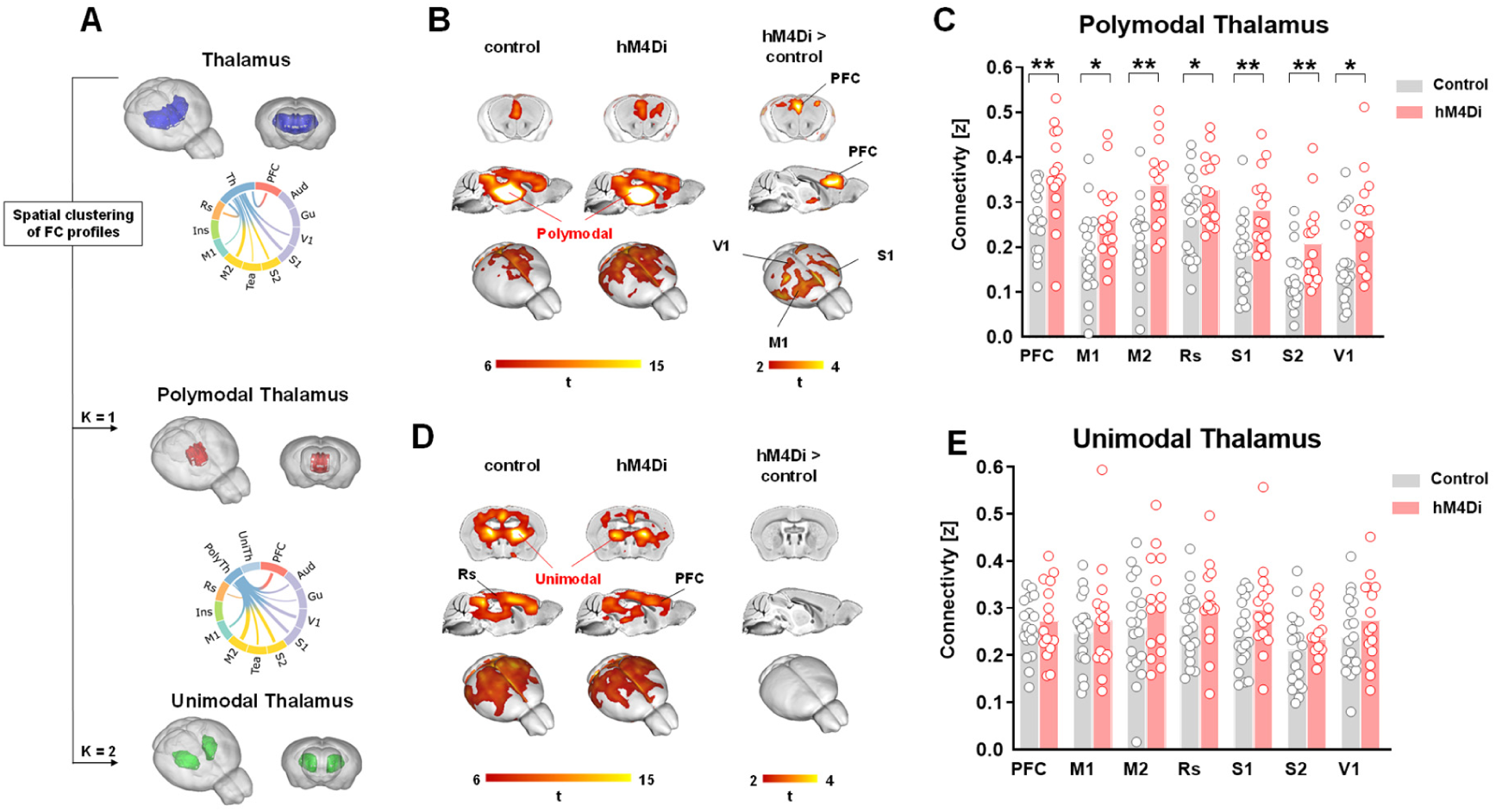
The polymodal thalamus is functionally overconnected to cortical areas. (A) k-means clustering of PFC-thalamic rsfMRI connectivity profiles (thalamus, blue; unimodal thalamus, green; polymodal thalamus, red) and circular representation of between-group connectivity difference of the entire thalamus and thalamic partition (UniTh: unimodal; PolyTh: polymodal thalamus). (B-D) Seed connectivity of sub-thalamic partitions, between group difference maps (B,C polymodal thalamus; D-E, unimodal thalamus) and corresponding quantification of thalamo-cortical connectivity (*p<0.05, **p<0.01; Benjamini-Hochberg corrected).

Notably, seed-based probing of the unimodal thalamus did not reveal significant differences between hM4Di-expressing and control animals (Z> 2.03, all voxels, cluster corrected at t > 2.03), whereas seed-based probing of the polymodal thalamus revealed in in hM4Di expressing-mice a widespread pattern of cortical overconnectivity, exceeding the boundaries of the PFC to encompass motor and somatosensory territories, including the retrosplenial cortex, as predicted by delta coherence measurements between these areas (Fig. 4; Z> 2.03, cluster corrected, p < 0.05, Fig. 5C, p <0.04 all regions, t test, FDR corrected, cortical ROI location in Fig. S10). To further probe the anatomical extension and specificity of these effects, we also mapped rsfMRI whole-brain connectivity in hM4Di and control animals using a brain-wide parcellation scheme (Figure S9). Corroborating our findings, this analysis revealed the presence of a significant cluster of overconnectivity in the DMN and in polymodal thalamic areas of DREADD-expressing animals, the latter region exhibiting overconnectivity with several cortical districts. No meaningful clusters of rsfMRI over-connectivity or under-connectivity were observed in any of the area examined. These findings are consistent with a a possible involvement of polymodal thalamic areas as putative propagators of delta activity and functional overconnectivity to cortical regions.

**Figure 5.**
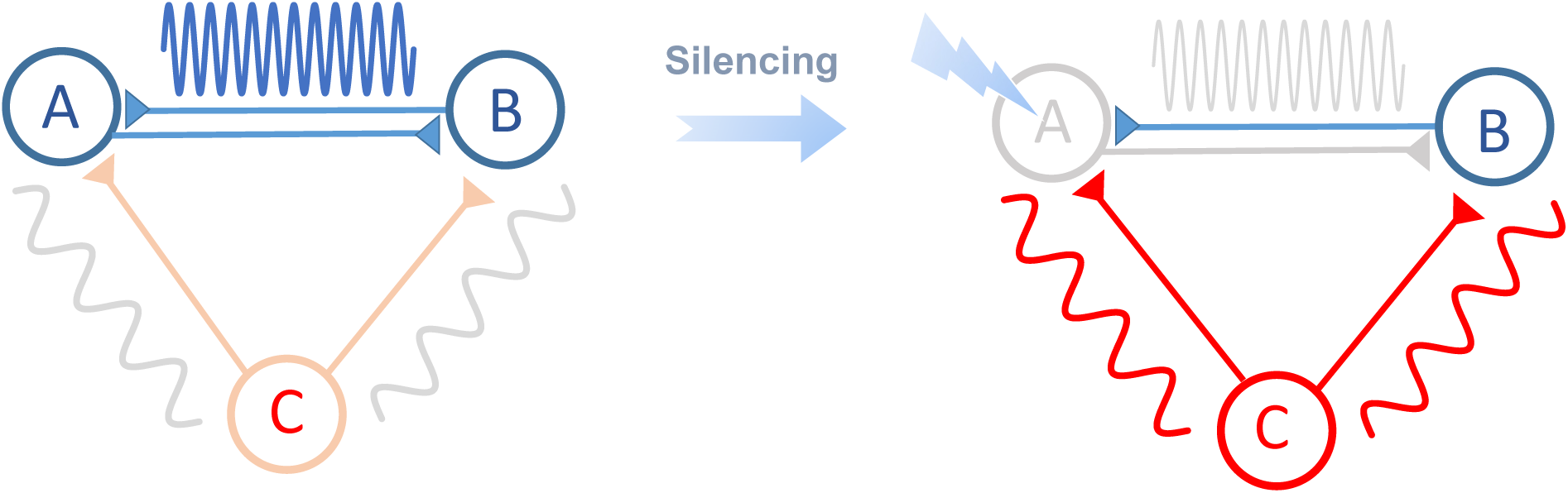
A possible interpretative model of our findings. Silencing cortical node A (i.e. PFC) reduces high frequency direct interactions between reciprocally innervated areas (A - B). The resulting functional overconnectivity reflects remotely generated (C) covarying slow oscillatory activity from brain rhythm generators (e.g., the polymodal thalamus, neuromodulatory nuclei, etc.). Slow frequency oscillatory activity may be further gated and propagated by polymodal thalamic nuclei to wider cortical areas.

Collectively, our findings support a putative interpretative model (sketched in Fig. 5) in which silencing of a cortical region (region A in Fig. 5) leads to reduced direct high-frequency axonal communication with its targets (area B in Fig. 5) and a concomitant increase in co-varying low frequency fluctuations, reflecting remote common input from slow rhythm generators (area C). Within this framework, interareal rsfMRI connectivity reflects captures both direct interareal communication, and common covariations due to large-scale synchronization. The paradoxical increase in rsfMRI connectivity between the silenced area and its targets would thus reflect a shift in the relative contribution of global low-frequency interactions with respect to direct higher-frequency rhythms. Our thalamic rsfMRI probing suggests that this effect may be possibly gated and propagated by polymodal thalamic nuclei to produce wider cortical over-connectivity.

## Discussion

Here we used neural perturbations to causally deconstruct brain-wide measurements of interareal functional connectivity based on rsfMRI. We found that chronic and acute neural silencing of the mouse PFC results in patterns of paradoxically increased rsfMRI connectivity with terminals directly innervated by the affected area. Further electrophysiological investigations revealed that chemogenetic silencing of the mouse PFC is associated with reduced gamma activity, and prominently increased interareal delta coherence, arguing against a vascular origin for the observed hyper-connectivity, and implicating a key contribution of slow oscillatory rhythms to the establishment of brain-wide rsfMRI coupling.

The observation that structural and functional connectivity strength are correlated both at the whole-brain and mesoscopic scale^4-6^ lies at the basis of structurally-based models of rsfMRI connectivity, according to which spatiotemporal correlations in spontaneous rsfMRI activity are critically shaped and constrained by underlying axonal connectivity^10, 12, 37^. A key prediction of these models is that regional inactivation of a neural node would result in reduced functional synchronization in distributed target areas, the spatial extent of which is dictated by the structural topography of the affected region^12^. Leveraging the recent implementations of chemo-fMRI in the mouse^20^, here we causally probed how cortical silencing affects brain-wide rsfMRI coupling. Motivated by reports describing hyperconnectivity in clinical conditions characterized by physical disruption of critical nodes^15, 16^, we first carried out proof-of-principle experiments using chronic silencing of neural activity in the mouse PFC via overexpression of a potassium channel. We next ran a set of mechanistic investigations leveraging acute chemogenetic inactivation of prefrontal activity via DREADDs. Surprisingly, we found that both chronic and acute silencing of the PFC paradoxically increased rsfMRI connectivity within thalamo-cortical substrates of the mouse DMN, encompassing cortical and subcortical regions densely innervated by prefrontal areas.

Our results advance our understanding of the principles underlying brain-wide rsfMRI coupling in two main directions. First, the observed breakdown between rsfMRI coupling and cortical activity challenge prevailing structurally based interpretations of functional connectivity, suggesting that rsfMRI-based functional connectivity does not monotonically reflect direct axonal output under all states and conditions, but can be critically modulated by remote sources of large-scale covariation (e.g. δ and infraslow oscillations^38, 39^), whose effect on functional connectivity can be as important as that of direct high frequency (e.g. γ) interareal interactions. It should be emphasized here that this result should not be intended as a refutation of the strong structural basis of rsfMRI connectivity, but rather the basis of an updated model in which large-scale cortical coupling is strongly biased by synchronized input from global rhythm generators. Second, our results also provide a novel reference framework for the interpretation and reverse engineering of rsfMRI overconnectivity observed in brain pathology, especially in degenerative or neurological states characterized by rsfMRI oversynchronization. Although compounded by pathophysiological rearrangements in synaptic activity ^40^, observations of unexpected increases in functional connectivity in neurological conditions characterized by loss of cortical function such as Alzheimer’s disease or stroke have been speculatively conceptualized as the result of compensatory rerouting of signal propagation along indirect structural paths, a neuroadaptive strategy aimed at maintaining task performance^15, 16, 41^. Our findings offer an alternative, network-level mechanism for these clinical observations, suggesting that rsfMRI overconnectivity may reflect a shifted balance between the amount of interareal synchronization due to direct axonal communication and low-frequency brain-wide synchronization mediated by single centers such as the polymodal thalamus ^30^ or by more global modulatory input ^42, 43^, a conceivable scenario in early-stage degenerative or neurological states characterized by loss of neuronal function in higher order cortical areas. EEG and MEG studies in stroke patients and neurodegenerative brain disorders provide indirect support to this hypothesis, as rsfMRI overconnectivity and robust δ hyper-synchronization has been observed ipsilaterally to lesioned areas in stroke ^16, 44^, and in Alzheimer’s disease patients in which this effect appears to be especially prominent in polymodal cortical areas^45, 46^. Within such a framework, functional impairments in cortical activity following degenerative pathology could manifest as hyper-synchronization during pre-disease states, eventually reverting to hypotrophy-associated under-connectivity at advanced stages of brain pathology.

While a neural mass phenomenon like interareal rsfMRI coupling cannot be mechanistically dissected into canonical circuital elements, our observation of prominently increased δ band coherence upon silencing of the PFC is consistent with prior reports linking spontaneous fMRI activity to intrinsic slow oscillatory activity^34, 39, 47-51^, and putatively implicates of a number of global rhythm generators to the establishment of brain-wide coupling. The fact that full expression of δ activity requires thalamic participation^52, 53^, together with the critical role of higher-order thalamic nuclei in enhancing and relaying δ synchronization^30^ point at a possible indirect involvement of thalamus in the establishment of rsfMRI over-connectivity. However, several other modulatory mechanisms, or subcortical substrates such as the claustrum, ^54^ may similarly (or alternatively) contribute to the observed oversynchronization, including the back-propagation of cortically generated slow oscillatory activity^55^, as well as a role for nested or adjacent frequency bands^51, 56^.

Because light sedation enhances slow oscillatory rhythms^57^, the sign and neurophysiological signature of the resting state connectivity changes we map here could conceivably be affected by brain states and arousal level. Such a scenario does not however diminish the relevance of our findings for two reasons. First, our observation of DMN overconnectivity in the face of robust neural silencing is sufficient to disprove a general monotonic relationship between axonal connectivity and rsfMRI coupling, causally demonstrating a non-dyadic relationship between these two entities, with implications both for the mechanistic modelling of rsfMRI and the reverse engineering of overconnectivity in brain disorders. Second, the light sedation regimen employed in this study is physiologically stable, preserves sensorial responsivity ^58^ and does not induce burst-suppression or cortical hyper-synchronization ^59^, resulting in preserved rsfMRI connectivity topography ^6, 21^ and rich fMRI network dynamics ^17^. These properties make the specific state under which our recordings were carried out not dissimilar from the quiet wakefulness state, often bordering sleep^60^, that characterizes most rsfMRI scanning in humans ^43^. In keeping with this notion, intracranial recordings have revealed the presence of significant epochs of delta activity under quiet wakefulness in rodents^61, 62^ and humans ^63^, and state-dependent electrophysiological measurements in humans have consistently shown that slow (delta and infraslow) cortical potentials represent the most solid and reliable correlate of rsfMRI coupling in sleep as well as in awake states^39, 64^. Because of the difficulty in controlling restraint-induced stress in highly rousable rodent species^43, 65^, the state-dependence of our manipulation is an interesting subject for future investigations which may benefit from the implementation of hemodynamic mapping in quite wakefulness, for example using functional ultrasound imaging ^66^.

Several experimental factors may underscore the contrast between our findings and the results of Grayson et al. ^10^. These include the specific wiring diagram of the regions silenced, given that the PFC and the amygdala are ontologically segregable, and exhibit different interactions with subcortical rhythm generators. Further experimental discrepancies, including the lack of electrophysiological recordings and a direct control for the unspecific hemodynamic and neural effects ^20, 67^ of CNO *per se* in Greyson et al. prevent a direct comparison of our findings, and could explain these differences. Importantly, these authors carried out their investigations under light sedation, arguing against a direct contribution of state-dependent differenced in the sign and distribution of the rsfMRI changes observed here. We also note that two recent rodent studies reported weak rsfMRI suppression upon acute silencing of dorsal cingulate cortex in awake restrained rats, and in anesthetized mice respectively^68, 69^. Neither investigation however reported electrophysiological evidence of suppression of spontaneous activity, nor incorporated a direct control group for the unspecific effects of the employed ligands, again preventing a direct comparison with the results we describe here. The recent observation that DREADD-induced increased firing produces rsfMRI desynchronization ^70^ in sedated mice is instead in good agreement with our results and interpretative model, pointing a bidirectional relationship between cortical activity and interareal coupling.

In conclusion, our data show that cortical silencing leads to paradoxical brain-wide rsfMRI overconnectivity and increased interareal delta coherence. These findings challenge prevailing interpretations of functional connectivity as a monotonic index of direct axonal communication and highlight a critical contribution of slow rhythm generators to the establishment of brain-wide functional coupling. Our work also defines novel testable network-level mechanisms for reverse engineering rsfMRI overconnectivity in clinical conditions characterized by loss of cortical function.

## Acknowledgements

This work has received funding from the European Research Council (ERC) under the European Union’s Horizon 2020 research and innovation programme (#DISCONN; no. 802371 to A.Go.), the Brain and Behavior Foundation (NARSAD; Independent Investigator Grant; no. 25861) and Simons Foundation (602849 to S.P). A.Go. also acknowledges support by the Simons Foundation (SFARI 400101), the NIH (1R21MH116473-01A1) and the Telethon foundation (GGP19177). We thank Michael Lombardo, Elizabeth De Guzman and all the members of the Gozzi laboratory for critically reading the manuscript. We also thank Thomas Gozzi for figure renderings.

## Author contribution

A.Go. conceived and supervised the study. CC carried out viral transduction and the fMRI experiments with help from A.Ga. FR designed, carried out and analyzed electrophysiological recordings. GI, SP, SV, NS, FR, CC, DGB and A.Go. analyzed rsfMRI and electrophysiological data. A.Go. and S.P. secured funding. A.G. and S.P. wrote the manuscript

## Experimental Procedures

### Ethical statement

All in vivo experiments were conducted in accordance with the Italian law (DL 26/214, EU 63/2010, Ministero della Sanità, Roma). Animal research protocols were reviewed and consented by the animal care committee of the University of Trento and Italian Ministry of Health (authorization no. 852/17 to A.G.). All surgical procedures were performed under anesthesia.

### Animals

Adult (6 week old) male C57Bl6/J were purchased from Jackson Laboratories (Bar Harbor, ME, USA). Mice were housed with temperature maintained at 21 ± 1°C and humidity at 60 ± 10%.

### Anatomical definition of mouse medial prefrontal cortex

Our anatomical definition of mouse medial prefrontal cortex (PFC) reflects recent neuroanatomical ^71^ and cytoarchitectural cross-species comparisons ^72^, according to which the mouse PFC comprises a prelimbic region, corresponding to primate Brodmann area 32 (A32), the anterior cingulate cortex, corresponding to Brodmann area A24b, and the infralimbic cortex, corresponding to Brodmann area A24a. Our viral manipulations were therefore aimed to silence an anatomical ensemble comprising all the above mentioned regions at the following coordinates, expressed in millimeter from Bregma: 1.7 from anterior to posterior, +/−0.3 lateral, −1.7 deep ^73^.

### Viral injections

Mice were anesthetized with isoflurane and head-fixed in a stereotaxic apparatus (Stoelting). Injections were performed with a Hamilton syringe mounted on Nanoliter Syringe Pump with controller (KD Scientific), at a speed of 0.05 μl/min, followed by a 5–10 min waiting period, to avoid backflow of viral solution. The following injections volumes were employed: 300 nL (AAV8-hSyn-hM4D(Gi)-mCherry and AAV8-hSyn-GFP; http://www.addgene.org) or 2 μL (AAV8-hSyn-MYC-mKir2.1(E224G/Y242F)-IRES-GFP, ^22^, http://www.vectorbiolabs.com or AAV8-hSyn-GFP; http://www.addgene.org) of viral suspension were injected bilaterally in the mouse PFC (see coordinates above). rsfMRI or electrophysiological recordings were carried out no sooner than three weeks after the injection, to allow for maximal viral expression.

### rsfMRI acquisitions

The animal preparation protocol was recently described in detail ^17, 74, 75^. Briefly, mice were anesthetized with isoflurane (5%, induction), intubated and artificially ventilated (2%, surgery). The left femoral artery was cannulated for continuous blood pressure monitoring. At the end of surgery, isoflurane was discontinued and substituted with a shallow halothane regimen (0.75%) to obtain light sedation and to preserve cerebral blood flow auto-regulation ^76^. Ventilation parameters were adjusted to maintain normo-physiological p_a_CO_2_ (< 40 mmHg) and p_a_O_2_ levels (> 90 mmHg, corresponding to >98% hemoglobin saturation).

rsfMRI data acquisition commenced 30 min after isoflurane cessation. Functional images were acquired with a 7T MRI scanner (Bruker, Biospin) as previously described ^77^, using a 72 mm birdcage transmit coil and a 4-channel solenoid coil for signal reception. Single-shot BOLD rsfMRI time series were acquired using an EPI sequence with the following parameters: TR/TE 1000/15 ms, flip angle 60°, matrix 98 x 98, FOV 2.3 x 2.3 cm, 18 coronal slices, slice thickness 550 µm.

rsfMRI acquisition with Kir2.1-transduced (AAV8-hSyn-MYC-mKir2.1(E224G/Y242F)-IRES-GFP, n = 16) and control mice (AAV8-hSyn-GFP, n = 19) encompassed 35-minute long timeseries, corresponding to 2100 volumes.

Chemo-fMRI acquisitions comprised two consecutive rsfMRI timeseries, encompassing 1800 volumes (30 min) and 2100 volumes (35 min), respectively. CNO (2 mg/kg, Sigma Aldrich) was injected intravenously fifteen minutes (volume #900) after the start of the first scan. The first 900 fMRI volumes of this first timeseries scan were used as pre-CNO baseline rsfMRI reference in time-resolved analyses. Based on the pharmacokinetic profile of CNO, the post CNO window was split into temporal domains as follows: the first 15 min post injection (900 volumes) were considered part of a drug equilibration window, while the following 35 min (2100 volumes) were considered to cover the DREADD active time window ^27^. All group comparisons in the chemo-fMRI study were carried out within this latter time window, unless otherwise stated. After post-mortem analyses of viral expressions, a total of N = 15 hM4Di and N= 19 GFP-transduced animals were retained for analyses.

### Image preprocessing and analysis

Raw rsfMRI timeseries were preprocessed as described in previous work ^17, 74^. Briefly, the initial 120 volumes of the time series were removed to allow for gradient equilibration effects. Data were then despiked, motion corrected and spatially registered to a common reference template. Motion traces of head realignment parameters (3 translations + 3 rotations) and mean ventricular signal (corresponding to the averaged BOLD signal within a reference ventricular mask) were used as nuisance covariates and regressed out from each time course. All rsfMRI time series also underwent band-pass filtering within a frequency window of 0.01–0.1 Hz and spatial smoothing with a full width at half maximum of 0.6 mm. To control for the effects of global fMRI signal regression on the mapped changes, all rsfMRI timeseries were also recomputed by regressing average fMRI signal within an intracerebral mask

rsfMRI connectivity of the mouse DMN in Kir2.1 and chemo-fMRI scans was probed using a seed-based approach. In the case of the chemo-fMRI study, this quantification was carried out during the CNO active time window. A 5×5×2 seed region was selected to cover the PFC areas targeted by viral injections. Voxel-wise intergroup differences in seed-based mapping were assessed using a 2-tailed Student’s t-test (|t| > 2, p < 0.05) and family-wise error (FWE) cluster-corrected using a cluster threshold of p = 0.05 as implemented in FSL (https://fsl.fmrib.ox.ac.uk/fsl/). The antero-posterior connectivity profile of the DMN was assessed by computing Person correlation between the PFC seed abovementioned and a series of 6 x 6 x 2 voxel seeds placed along the midline extension of the cingulate and retrosplenial cortices as previously described ^25^. Quantification of cortico-thalamic connectivity was carried out with respect to a meta-regional parcellation of the mouse cortex in volumes-of-interest (VOIs), Fig. S10. To rule out a possible confounding contribution of spurious neurovascular changes in CNO-induced rsfMRI connectivity alterations, we calculated and statistically compared the characteristic hemodynamic response function between kir2.1 and control mice, and between hM4Di-expressing and control mice upon CNO-administration (active phase), as previously described ^25, 78^.

Whole-brain connectivity in hM4Di and Control mice was calculated across a set of volumes of interest recapitulating anatomical areas of the Allen brain atlas. The anatomical probed area were selected according to their coverage of previously characterized network systems of the mouse brain ^21, 74, 77^: TH: thalamus (thalamus polymodal association cortex related, Thalamus Sensory-Motor cortex related); STR: striatum (Striatum dorsal region left, striatum dorsal region right, striatum ventral region left, striatum ventral region right); LCN: lateral cortical network (LCN: primary motor cortex left, primary motor cortex right, primary somatosensory cortex left, primary somatosensory cortex right, secondary somatosensory cortex left, secondary somatosensory cortex right, Lateral septal complex left, lateral septal complex right); HCP: hippocampus (Ammon’s horn left; Ammon’s horn right, dentate gyrus left; dentate gyrus right, Entorhinal area left, entorhinal area right, subiculum left, subiculum right); DMN: default mode network (anterior cingulate area; Infralimbic area, secondary motor cortex left, secondary motor cortex right, orbital area, prelimbic area, posterior parietal association areas left, Posterior parietal association areas right, retrosplenial area); To relate the strength of underlying anatomical connectivity to the regions exhibiting increased rsfMRI connectivity with voxel resolution we extracted outgoing projections from the affected PFC regions using a spatially-resampled (0.027 mm^3^) version of a voxel scale model of the Allen Brain Institute structural connectome ^6^. We then plotted the strength of PFC-departing structural projections against the corresponding between-group difference in rsfMRI connectivity using the cluster-corrected difference map, and assessed differences in the distribution of overconnected areas with respect to all the brain voxels using a Wilcoxon rank-sum test.

To quantify the contribution of distinct thalamic subregions to overall group differences, we used k-means clustering to partition voxels within the thalamus, based on whole-brain rsfMRI group-difference obtained using the PFC seed as recently described (Arthur and Vassilvitskii, 2007; Schleifer *et al*, 2019). This approach revealed two major thalamic clusters, one medial and one bilateral partition encompassing sensory areas. Seed-based functional connectivity was subsequently computed for each of the two-resultant k-means clusters independently and the resulting functional connectivity maps compared and quantified across cortical VOIs.

### Electrophysiological recordings

Electrophysiological recordings were carried out in animals subjected to the same animal preparation and sedation regime employed for rsfMRI mapping (Ferrari *et al*, 2012; Sforazzini *et al*, 2014). Briefly, mice were anaesthetized with isoflurane (5% induction), intubated, artificially ventilated (2% maintenance), and head-fixed in a stereotaxic apparatus (Stoelting). The tail vein was cannulated for CNO injection. To ensure maximal consistency between viral injections and recording site, the skull surface was exposed and an insertion hole in the right PFC was gently drilled through the skull corresponding to the location of prior viral injection point. A single shank electrode (Neuronexus, USA, interelectrode spacing 1 - 2.5 mm) was next inserted through the overlying dura mater by a microdrive array system (Kopf Instruments, Germany) at an insertion rate of 1 µm/min to reach the same stereotaxic coordinates employed for viral injection.

The site receptive fields were plotted manually and the position and size of each field were stored together the acquisition data. After electrode insertion, isoflurane was discontinued and replaced by halothane at a maintenance level of 0.75%, to induce rsfMRI-comparable sedation. Electrophysiological data acquisition commenced 1 hour after isoflurane cessation. Such transition time was required to ensure complete washout of isoflurane anesthesia and avoid residual burst-suppressing activity associated with extended exposure to deep anesthetic levels.

Neural activity was next recorded in consecutive 5 min time bins to cover a 15 min pre-injection time window, and a 60 min post CNO time-frame in n = 5 hM4Di and n = 5 GFP-expressing mice. Signals were amplified using an RHD 2000 amplifier system (Intan Technologies) at a sampling rate of 20 kHz. In the case of control Kir2.1 recordings, a four shank electrode was inserted along the coronal plane to bi-hemispherically cover the right (Kir2.1-expressing) and left (GFP-expressing) PFC (n = 4). The left region served as internal reference control to better assess the efficacy of Kir2.1 silencing. Electrophysiological signals were then recorded into 5 min time bins to cover a 35 min time-window.

To measure multi-electrode coherence, three electrodes were inserted in key cortical and subcortical substrates identified as overconnected in our chemo-fMRI mapping. A multi-probe micromanipulator (New-Scale Technologies) was used to insert a 32 channels four shank electrode (Neuronexus, USA, interelectrode spacing 1 - 2.5 mm) in the right prefrontal cortex, and two 16 channels single shank electrode (Neuronexus, USA, interelectrode spacing 1 - 2.5 mm) in the right medio-dorsal thalamus and retrosplenial cortex, respectively (insertion location in Fig. S6). To reduce tissue damage, an insertion rate < 1 µm/min was employed, allowing for a 30 minute equilibration every 400 µm traveled. Ground electrodes were put in contact with the cerebral brain fluid through a window drilled in the skull. Signals were then recorded into 1 min time bins to cover a 15-min pre-injection baseline and a 40-min post CNO time window in n = 5 hM4Di and n = 5 GFP-expressing adult C57Bl6/J mice.

#### LFP and multi-unit activity (MUA)

To compute the LFP signal, raw extracellular recordings were first down-sampled to 4 kHz, then band-pass filtered to 1-250 Hz using a previously published two-step procedure ^79^. Briefly, raw timeseries were first low-pass filtered using a 4^th^ order Butterworth filter with cut-off frequency of 1 kHz. The resulting timeseries were next down-sampled to 2 kHz, then again filtered using a Kaiser window filter between 1Hz to 250Hz (with a sharp transition bandwidth of 1Hz, passband ripple of 0.01 dB and a stop band attenuation of 60 dB) and then resampled at 1 kHz. All the filtering was applied both in forward and backward temporal direction to avoid any phase transitions due to filtering.

To compute multi-unity activity (MUA) we again followed the procedure described in ^79^. Briefly, we computed a band-passed high-frequency signal using a 4^th^ order Butterworth filter with a <100 Hz cut off frequency, then band-pass filtered between 400 and 3000 Hz using a Kaiser window filter (with transition band of 50 Hz, stopband attenuation of 60 dB, and passband ripple of 0.01 dB). From the high frequency signal we detected spike times using a spike detection threshold corresponding to 4-times the median of the high frequency signal, then divided by 0.6745 as suggested in (Quiroga *et al*, 2004). Spikes were considered to be biologically plausible, and as such retained in these computations, only if occurring more than 1 ms apart. This method does not permit the isolation of single units. For the purpose of this study, however, recording the activity of a small group of neurons provided a sufficient (if not better) estimate of local spike rate changes.

To quantify effectiveness of Kir2.*1* in suppressing spontaneous activity, firing rate was computed (in units of spikes/s) by dividing the number of spikes per electrode by the recording duration in seconds. The resulting spike rates were averaged across the channels corresponding to the virally targeted or control region. The average spike rate for four subjects were next tested against each other using paired student t test. To assess the effect of chemo-fMRI manipulations, spiking activity was computed in hM4Di and GFP-expressing mice as described above and segmented into one-minute bins. We next computed the channel averaged firing rate for each segment, and normalized this value with respect to the firing rate recorded during baseline (pre-CNO) period. The resulting baseline normalized firing rate index was then used to assess changes in spiking rate upon CNO injection in the two experimental cohorts.

To determine the time lag at which different samples from the same channels could be considered as approximately statistically independent we computed for each subject the autocorrelation of the channel-averaged firing rate for each subject, and computed the time lag necessary for the autocorrelation function to drop below a (95%) confidence interval. We then, for each subject, retained samples of the baseline-normalized firing rate at different times separated by the above obtained lags. We next separately analyzed the baseline-normalized rates in three different periods: 0 to 15 minutes (transient time), 15 to 35 minutes and 35 to 55 minutes after CNO the injection (active times). We pooled all the retained data points in these windows both over time and over subjects and then compared the median between the two populations using two-sided Wilcoxon rank-sum test. The obtained p-values were corrected using Bonferroni-Holm (1979) method for multiple comparisons correction.

LFP spectrograms were computed using a Fourier transform with a Kaiser window with a frequency resolution of 0.15 Hz, temporal resolution of approximately 6 seconds, and with 50% overlapping windows. Spectrograms and their differences were smoothed in time with the resolution of half a minute, and in frequency with the resolution of 1Hz using median filter. To quantify the effect of CNO on LFP rhythms, we computed a spectrogram modulation index as follows. First, the channel-averaged spectrograms for the duration of the baseline recording was computed. Next, we averaged time-frequency spectral profiles over time, resulting in frequency-resolved spectral profiles. The effect of CNO was assessed by computing a modulation index, defined as the ratio of channel-averaged spectrogram after injection minus baseline, time-averaged spectrogram and the sum of the same quantities, for every time window and for every frequency. This modulation index ranges between −1 and 1, and describes the changes due to drug injection over time, for each frequency.

To obtain a statistical assessment of CNO effects across groups and bands, we computed the autocorrelation of spectrograms for every subject at every frequency, and resampled the spectrograms using only uncorrelated data bins based on the lag times identified. We next computed the median of the modulation index over different frequency bands defined as follows: δ (0.1-4 Hz), θ (4-8 Hz), α (8-12 Hz), β (12-30 Hz), γ (30-70 Hz). Data within each band were pooled over uncorrelated time points (determined as above by taking time samples separated by lags at which autocorrelation became negligible) and over subjects, and the population medians were compared using two-sided Wilcoxon rank-sum tests, followed by Bonferroni-Holm correction.

### Multielectrode coherence

To analyze multi-electrode recordings, channel averaged spectrograms were preprocessed as described above. Temporal smoothing was carried out using a 60-second median filter as described above. CNO spectrograms (30-40 post CNO injection window), were normalized with respect to the last 3 minutes of pre-CNO baseline using a modulation index as before. To obtain a quantifiable assessment of the CNO effect across different groups and bands, data within each band were pooled into 60-second bins, and subject and the population medians were compared for each region separately using a two-sided Wilcoxon rank-sum tests, followed by Bonferroni-Holm correction. The use of 60 second bins was motivated by estimations of the time lags at which autocorrelation of electrophysiological signals become negligible (and thus samples are approximately independent) as described above.

To measure spectral power coherency, the magnitude of squared coherency was computed using Welch’s overlapped averaged periodogram method ^80^ with a 50% overlapping window of 2 seconds length. The coherency was calculated for every 60-second bin of channel-averaged recordings. To assess the effect of chemogenetic manipulations, the magnitude squared coherency for the 30 minutes post injection time was normalized with respect to the time averaged coherency recording during last 3 minutes of pre-CNO baseline at every time and frequency using a modulation index as described above. The obtained coherency modulation indices were next pooled into 60 second bins (containing uncorrelated data points), and the median over different frequency bands was calculated for each region pair and compared using two-sided Wilcoxon rank-sum tests, followed by Bonferroni-Holm correction.

To quantify interregional phase coupling, LFP data were first filtered using a third order Butterworth filter in delta band, and the instantaneous phase of each channel was computed by taking the phase of the analytical signal resulted from the Hilbert transform. For all possible pair of channels belonging to two different regions, we next computed the corresponding Phase Locking Value (PLV) as follows:

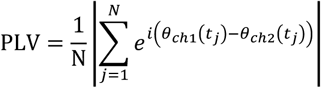

Where N is the number of data points in time and *θ*_*chl*_(*t*_*j*_), *θ*_*ch2*_(*t*_*j*_) are the instantaneous phase of the LFP of channel 1 and 2 at time j. The PLV value for each pair during the 30 to 40 minutes post CNO was next normalized with respect to the PLV value of the last 3 minutes of the pre-CNO baseline using a modulation index and was pooled over channel pairs and animals. The obtained population medians of the control and experimental group for each region pair were next compared using a two-sided Wilcoxon rank-sum tests, followed by Bonferroni-Holm correction.

## Supplementary Figures

**Figure S1.**
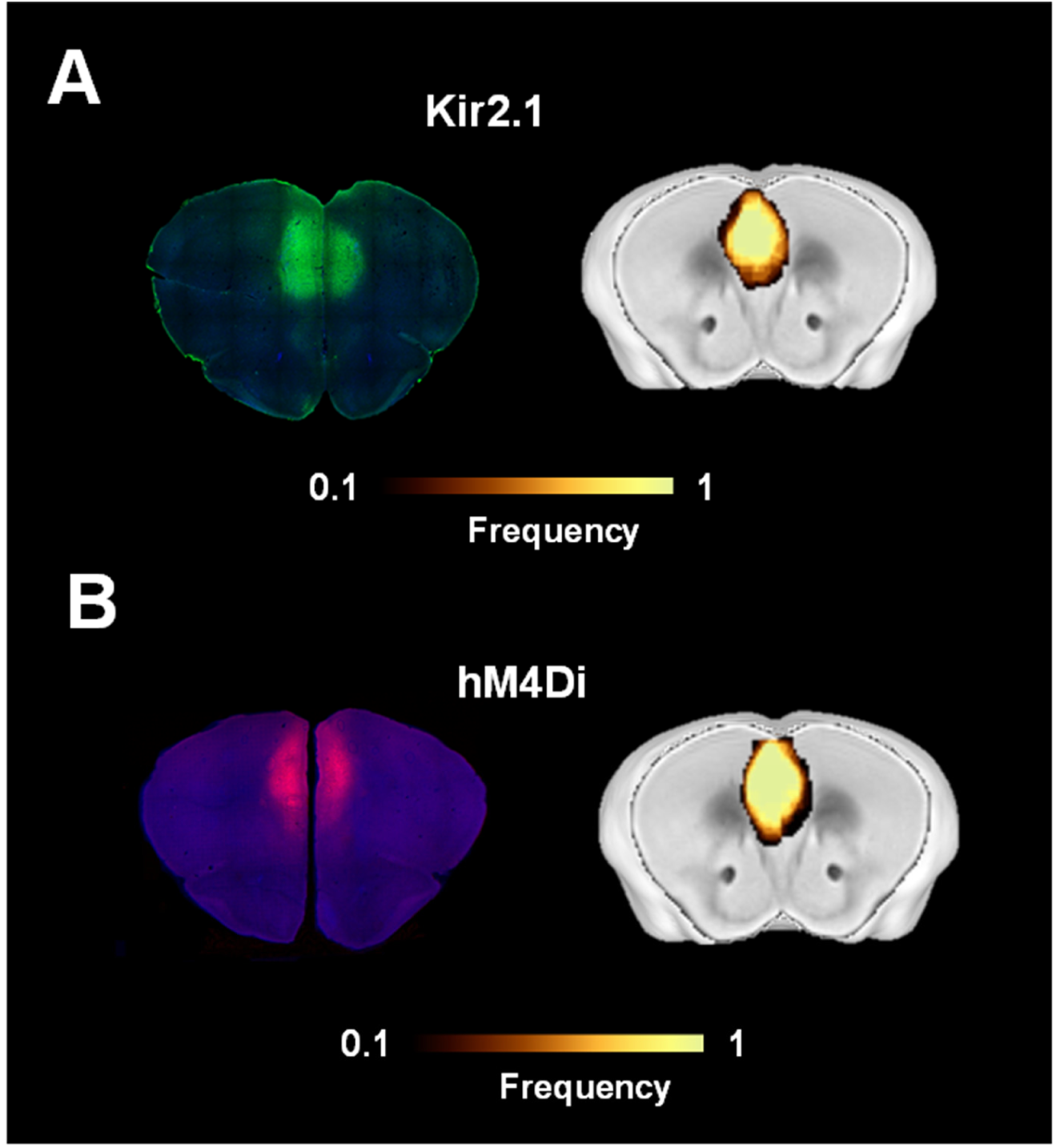
Viral expression localization. Brain sections showing (A) Kir2.1 (green) and (B) hM4Di (red) expression. Heat maps illustrate a qualitative regional assessment of viral expression across subjects.

**Figure S2.**
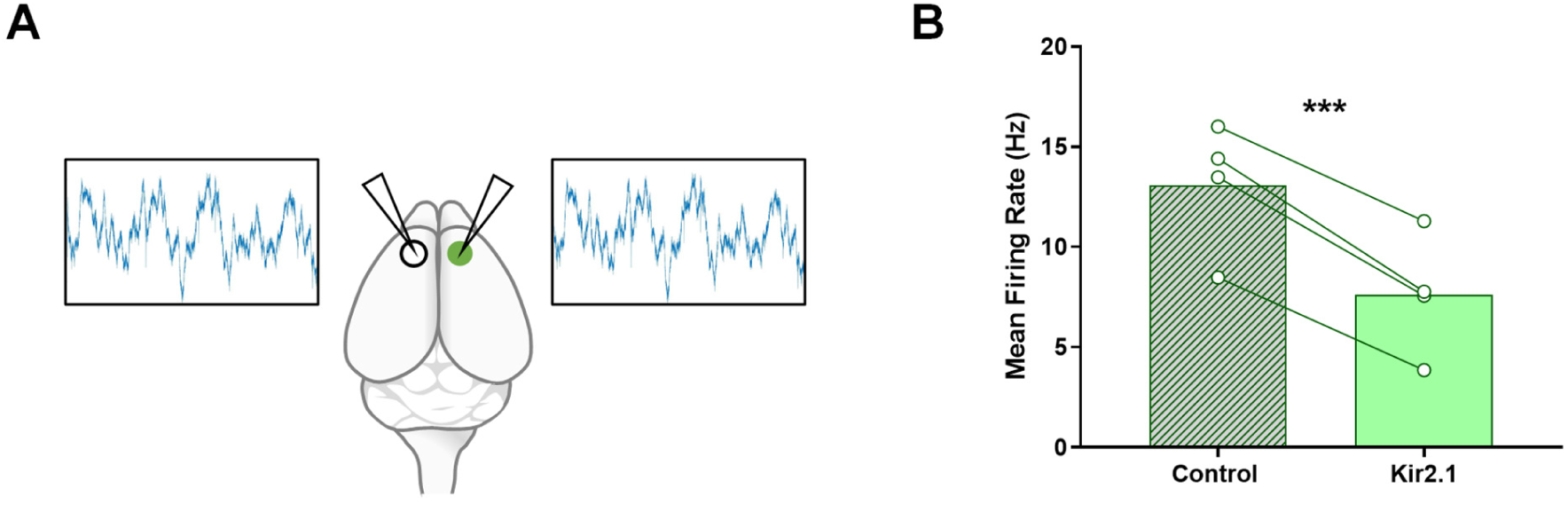
Overexpression of the potassium channel Kir2.1 in the PFC reduces spontaneous neural activity. (A) Experimental design: Kir2.1 injection was performed unilaterally in the right PFC. A viral vector encoding GFP was injected in the contralateral area. Electrophysiological recordings were carried out bilaterally using a four-shank electrode. (B) Mean spontaneous firing rate for the control side (no Kir2.1 expression), and the side expressing Kir2.1. (n=4; ** p<0.005).

**Figure S3.**
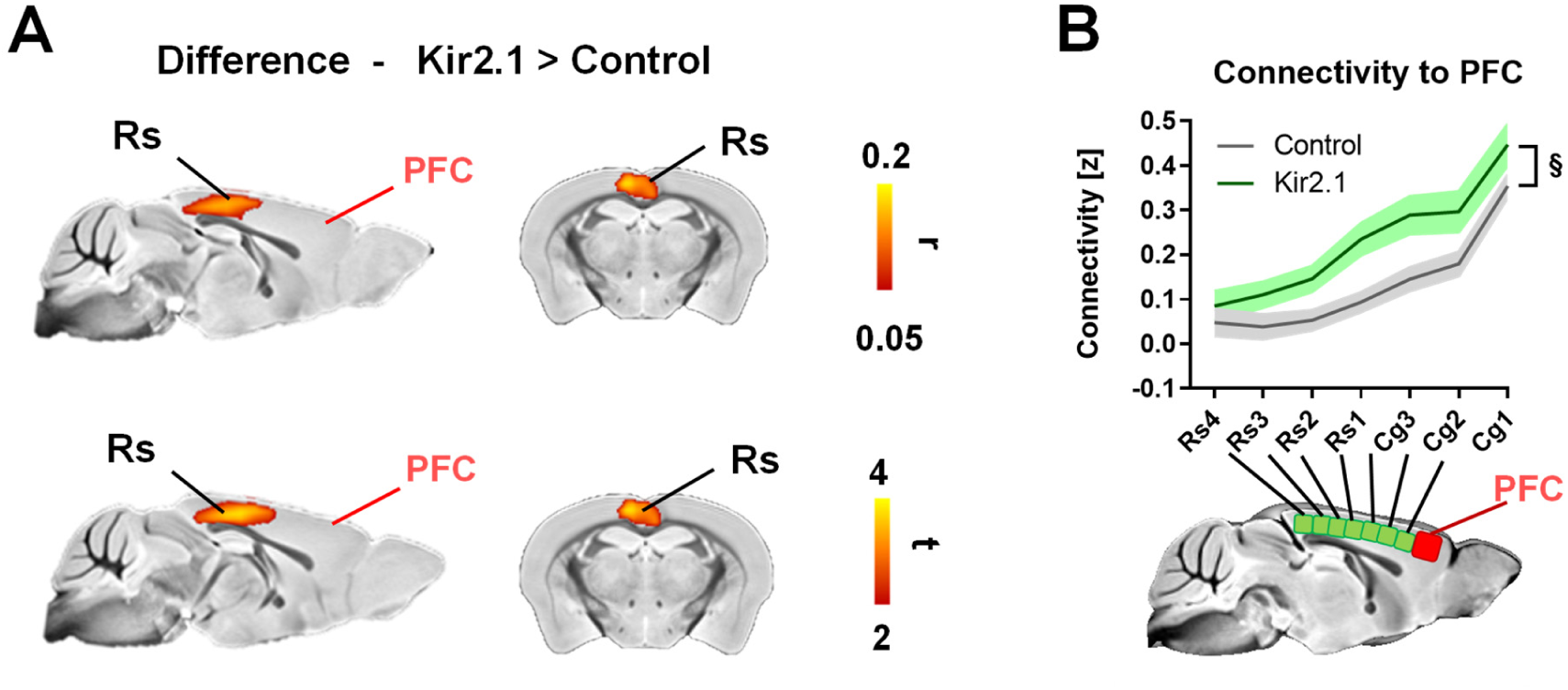
Foci of rsfMRI overconnectivity in Kir2.1 mice upon rsfMRI global signal regression (A) Between-group difference maps (Pearson’s r, and corresponding T stat difference maps). (B) Antero-posterior profiling of rsfMRI connectivity of the PFC along the midline axis of the mouse DMN in the two cohorts revealing consistent overconnectivity in Kir2.1 mice. rsfMRI connectivity was here computed after fMRI global signal regressions. Data are plotted as mean ± SEM. Cg: cingulate cortex; PFC: prefrontal cortex, RS: retrosplenial cortex; §p<0.05, 2-way ANOVA repeated measurements, genotype effect.

**Figure S4.**
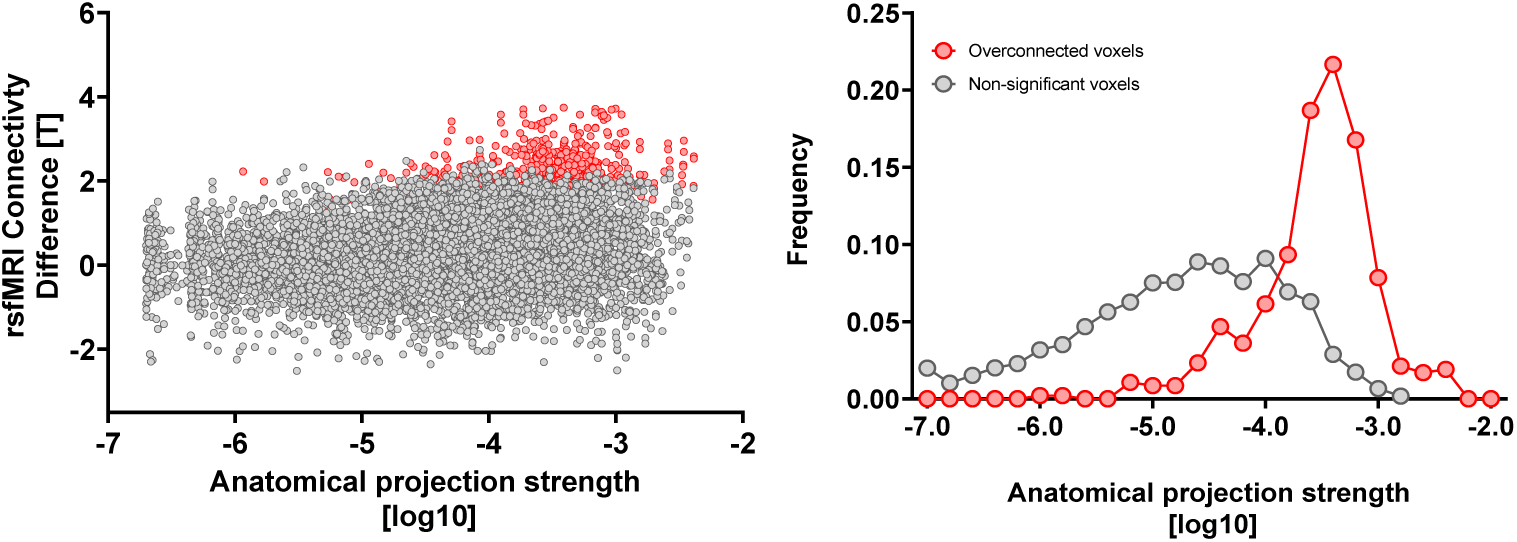
Voxels exhibiting rsfMRI overconnectivity upon chemogenetic silencing of the PFC are robustly innervated by the PFC. Left: scatter plot illustrating intergroup differences in rsfMRI connectivity as a function of PFC structural connectivity strength. Note that all significantly overconnected voxels (red) contain robust axonal projections from the PFC. Right: distribution of voxels exhibiting the most significant rsfMRI connectivity (red) and those that are not affected (grey).

**Figure S5.**
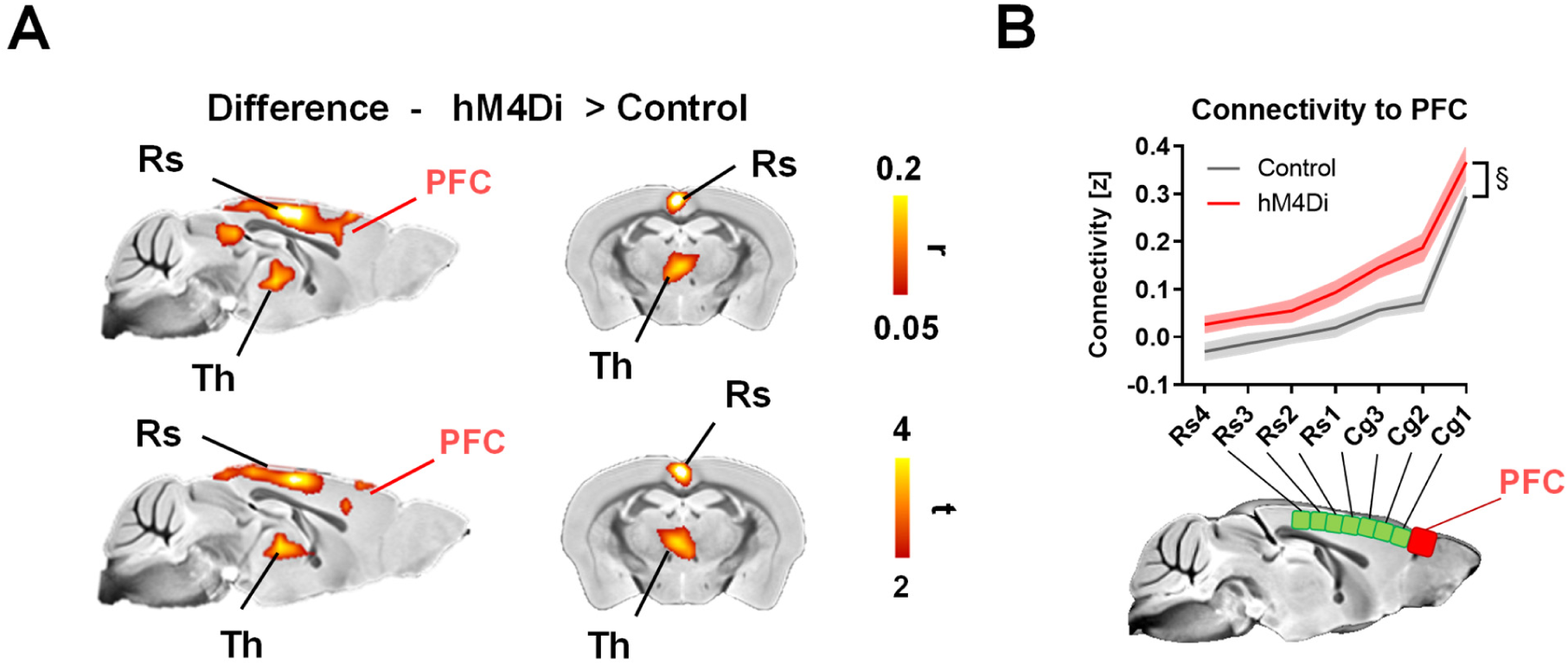
rsfMRI overconnectivity in hM4Di mice is not affected by rsfMRI global signal regression (A) Between-group difference maps (Pearson’s r, and corresponding T stat difference maps). (B) Antero-posterior profiling of rsfMRI connectivity of the PFC along the midline axis of the mouse DMN in the two cohorts. rsfMRI connectivity was here computed upon fMRI global signal regressions. Data are plotted as mean ± SEM. Cg: cingulate cortex; PFC: prefrontal cortex, RS: retrosplenial cortex; § p<0.05, 2-way ANOVA repeated measurements, genotype effect.

**Figure S6.**
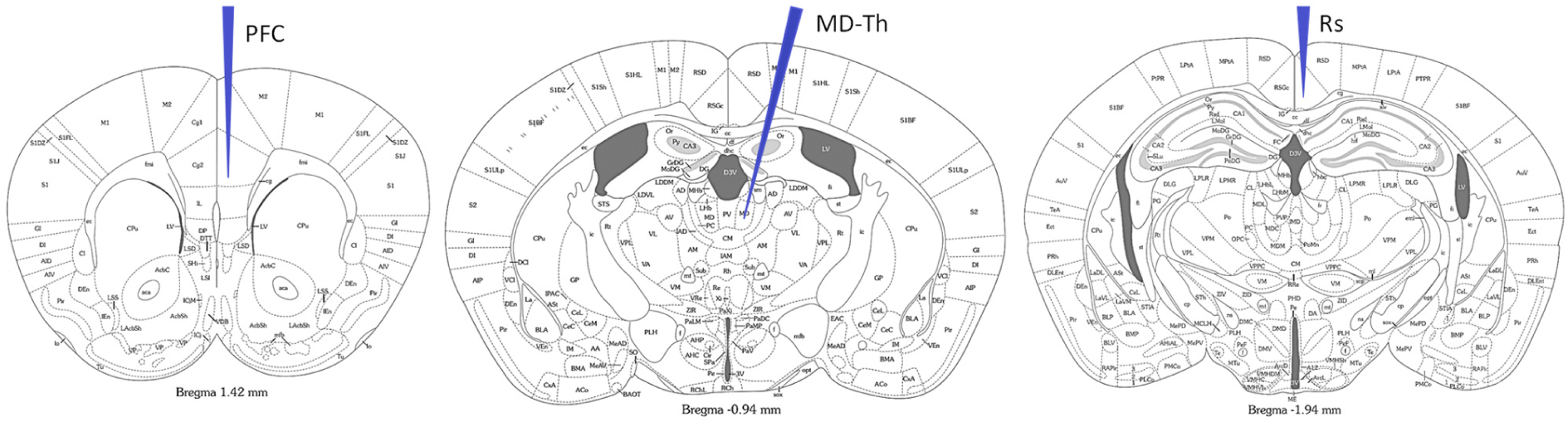
Anatomical location of electrode insertion site for multi-electrode neural recordings.

**Figure S7.**
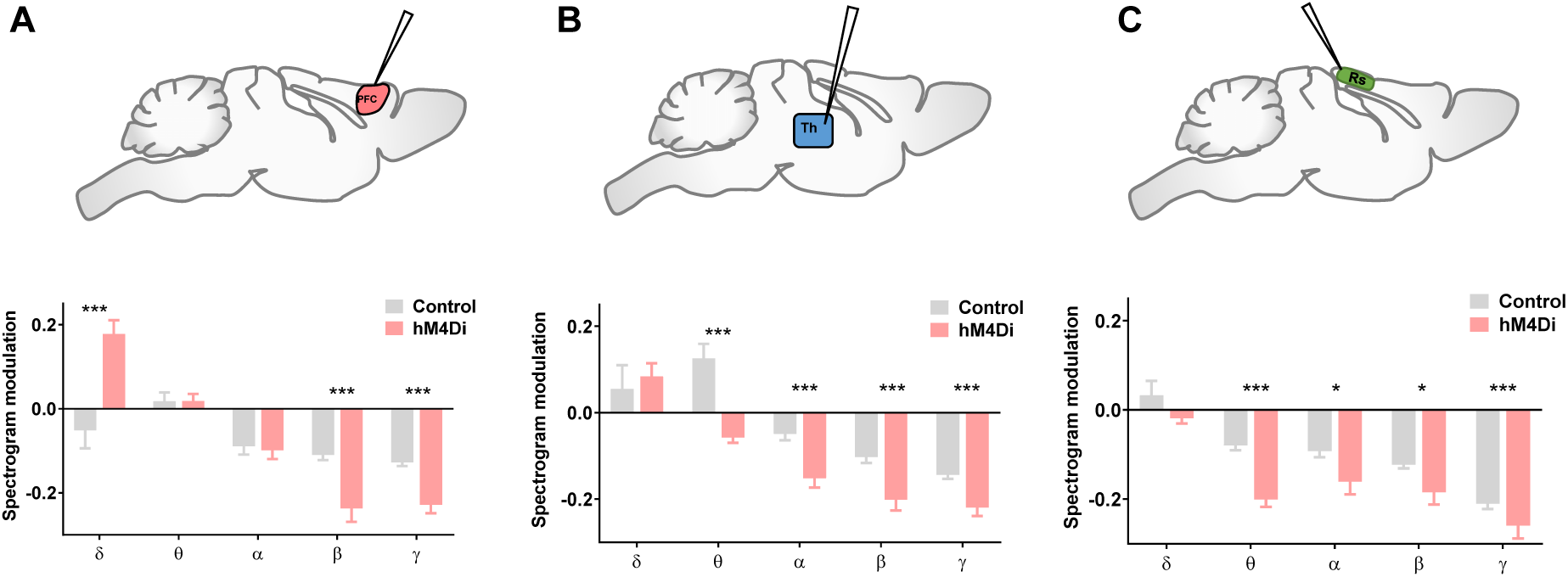
Quantification of band-specific power spectrum changes in LFPs recorded in the PFC (left), centromedial thalamus (Th; center) and retrosplenial cortex (Rs; right) upon systemic CNO administration. Power was quantified with respect to pre-injection baseline (mean ±SEM; *p <0.05, **p<0.01, ***p<0.001, t test, FDR corrected).

**Figure S8.**
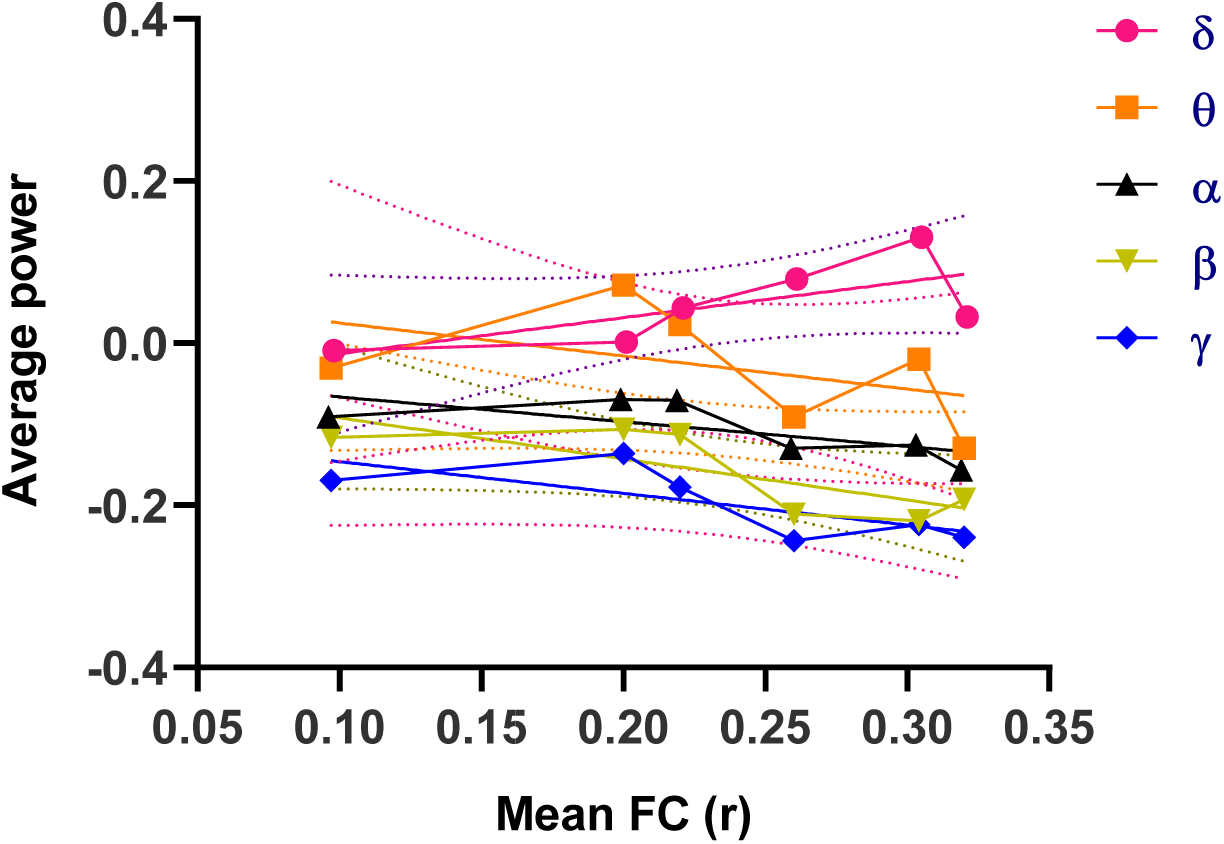
Correlation between mean band-specific power and corresponding mean functional connectivity for all pairs of electrophysiologically-probed regions (PFC-Rs; PFC-Th; Rs-Th). Mean FC and band-specific power were extracted for corresponding regional pairs during the CNO active time window in hM4Di and control animals.

**Figure S9.**
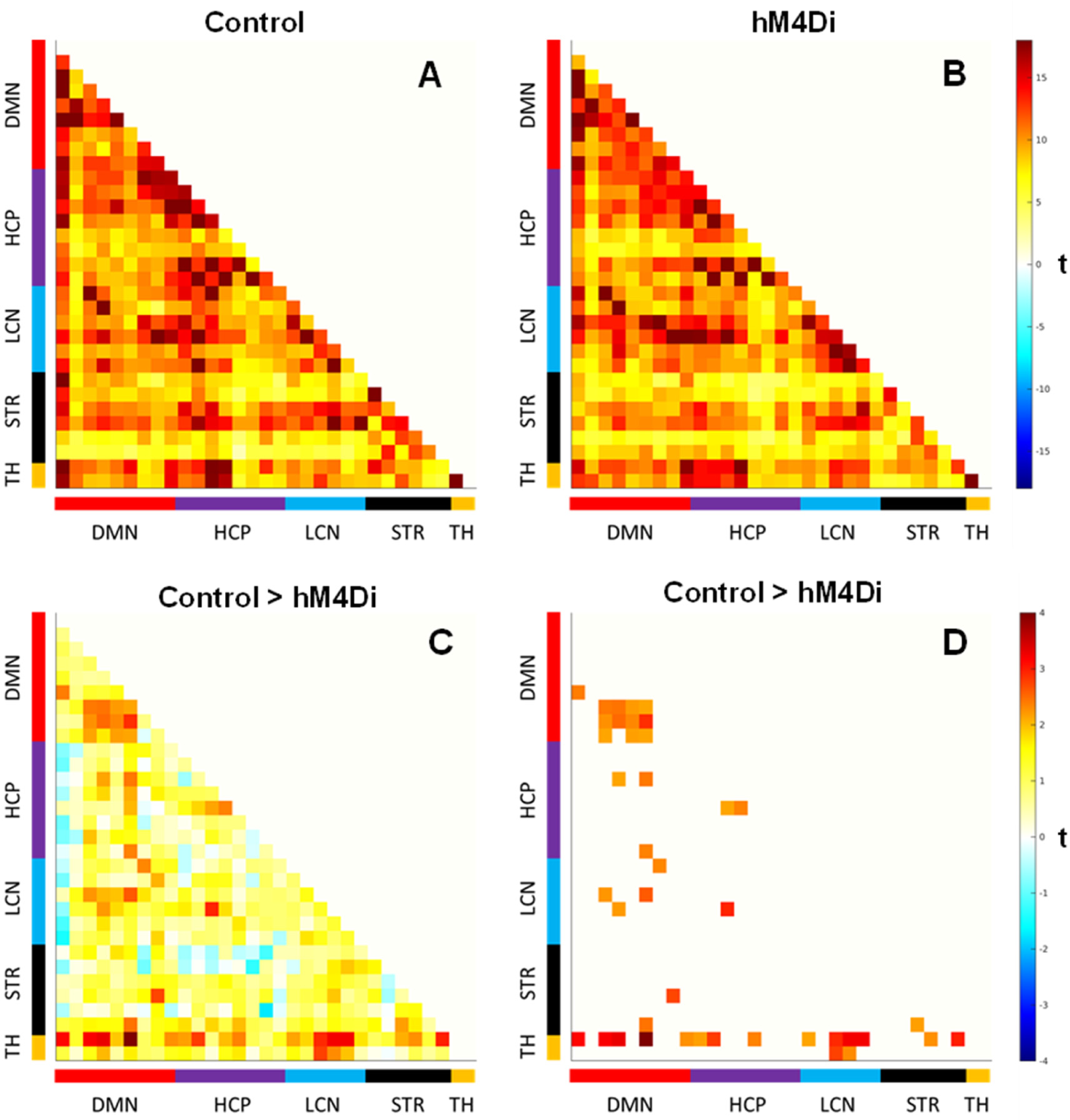
Whole-brain rsfMRI connectivity in control and HM4Di mice. Correlation matrices in (A) and (B) depict inter-areal connectivity in control and hM4Di mice respectively. (C) Mean difference map (t), and (D) regions exhibiting connectivity differences larger than |t|> 2.1, corresponding to p<0.02 two tailed (family-wise uncorrected). TH: thalamus; STR: striatum; LCN: lateral cortical network; HCP: hippocampus; DMN: default mode network.

**Figure S10.**
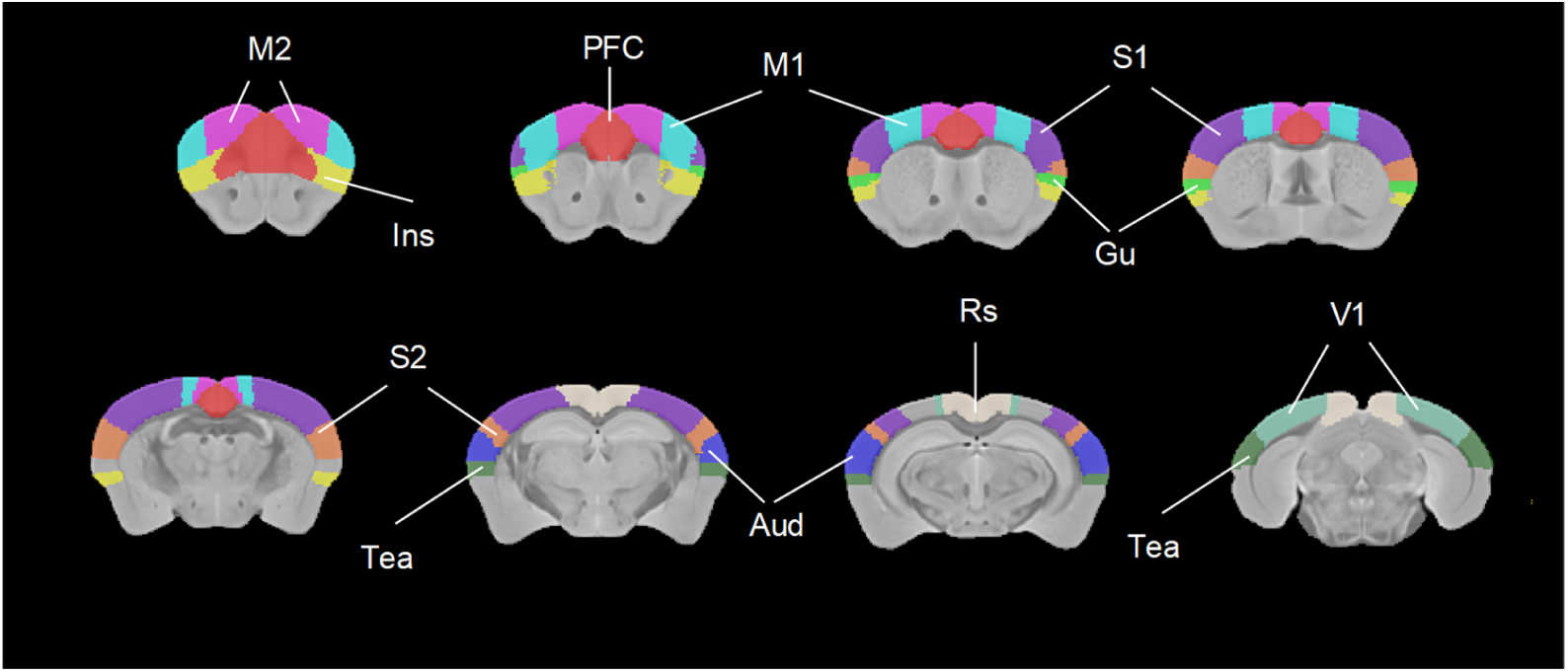
Anatomical location of volumes of interest employed for regional quantification of cortico-thalamic connectivity. Aud: auditory cortex; Gu: gustatory cortex; Ins: Insula; M1: primary motor cortex; M2: secondary motor cortex; PFC: prefrontal cortex, RS: retrosplenial; S1: primary somatosensory cortex; S2: secondary somatosensory cortex Tea: Temporal associate cortex; V1: primary visual cortex.

